# Pancreatic endocrine cells are transduced by adeno-associated virus serotypes 2 and 9 but not 6

**DOI:** 10.1101/2024.08.22.609291

**Authors:** Vishal Ahuja, Sanjeev Jeyabalan, Emmanuel S. Tzanakakis

**Affiliations:** Department of Chemical and Biological Engineering, Tufts University, Medford, MA 02155; Department of Developmental, Molecular and Cell Biology, Tufts University School of Medicine, Boston, MA 02111; Graduate Program in Pharmacology and Experimental Therapeutics and Pharmacology and Drug Development, Tufts University School of Medicine, Boston, MA 02111; Clinical and Translational Science Institute, Tufts Medical Center, Boston, MA 02111

**Keywords:** beta-cells, alpha-cells, pancreas, adeno-associated virus, infection, gene delivery

## Abstract

Adeno-associated viruses (AAVs) have emerged as powerful tools for delivery of genes to a variety of cell types including pancreatic endocrine cells. Currently, AAV serotype 8 (AAV8) is the main AAV vector employed for infecting pancreatic cells for transgene transfer. We aimed to address whether alternative serotypes (AAV2, AAV6, and AAV9) commonly used for gene transfer can be effective in transducing pancreatic cells efficiently. We also screened the additives heparin and neuraminidase to further understand the interaction between the individual AAV types included in this work and the cells for optimal infection. Murine pancreatic β-cells and α-cells as well as fibroblasts were infected with AAV serotypes 2, 6, and 9 carrying the transgene for enhanced green fluorescent protein (eGFP). AAV2 outperformed AAV9 in transducing pancreatic cells, while AAV6 induced cytotoxicity. Both AAV2 and AAV9 displayed slightly higher tropism for α-cells than for β-cells. Compared to the pancreatic cells, the fraction of GFP-expressing cells at various multiplicities of infection was consistently lower for fibroblasts. Incubation of AAV2 with heparin prior to transduction failed to induce any GFP expression in β-cells, indicating that the primary site used for initial interaction with pancreatic cells are heparan sulfate proteoglycans. Treatment of β-cells with neuraminidase prior to AAV9 infection appeared to improve the number of GFP-positive cells, but the increase was not statistically significant. These findings expand the repertoire of available serotypes for AAV-mediated delivery of transgenes to pancreatic endocrine cells and may contribute to gene therapy strategies for pancreas pathologies.

## INTRODUCTION

Adeno-associated viruses (AAVs) are emerging as major vehicles for gene delivery to various therapeutically useful cell types. Among these are pancreatic endocrine cells, which play a central role in blood glucose homeostasis and their damage is intimately linked to diabetes. Cell therapies are in development aiming to provide diabetic patients with fully functional endocrine cells^1^ including insulin-producing cells derived from stem cells^2,3^. In this vein, gene transfer has the potential to reduce autoimmune destruction of β-cells in type 1 diabetes (T1D)^4^ or ameliorate insulin homeostasis in type 2 diabetes (T2D)^5,6^. Hence, vectors capable of safe and efficient delivery of transgenes imparting curtailed immunogenicity and/or enhanced insulin secretion to native or transplanted cells are highly desirable.

Lacking the viral genes necessary for replication, AAVs are considered non-pathogenic and are less immunogenic than other viral vectors^7,8^. Moreover, AAV infection results in integration or episomal expression of transgenes in target cells for long-term gene expression^9^. Additionally, there are several serotypes of AAVs affording a wide range of tropism. To this end, there have been limited reports on AAV serotypes which are suitable specifically for genetic engineering of pancreatic cells. Most relevant works entail AAV infections of pancreatic tissue in rodents, particularly utilizing AAV8. For instance, glucose-stimulated insulin secretion (GSIS) is enhanced *in vivo* by delivering the genes encoding microRNA-132 under a rat insulin promoter using AAV8 for overexpression in β-cells^10^. Others reported improved β-cell persistence by overexpressing regulatory genes following transduction with AAV8^11,12^. Additionally, improved AAV8 infectivity of pancreatic cells has been achieved by introducing point mutations in the virus capsid^13^. AAV8 has also been used to identify rodent α-cells *in vivo* by introducing the enhanced green fluorescent protein (eGFP) transgene under the rat glucagon promoter^14^, indicating AAV8’s affinity for multiple endocrine pancreatic cell types.

Yet, the transduction efficiency of pancreatic cells by other common AAV serotypes such as AAV2, AAV6 and AAV9, remains unclear. AAVs interact with cells primarily by binding to glycans on their surface^15^. The primary binding sites for AAV2 are heparan sulfate proteoglycans (HSPGs)^16^. Prior studies have confirmed the presence of HSPGs on murine^17,18^ and human β-cells^19,20^, indicating the potential for AAV2 binding. In this case, the interaction of AAV2 with β-cells may be competitively inhibited by free heparin^21^, although relevant reports are conflicting and suggest that heparin can enhance the infectivity of other AAV serotypes^22^. AAV6 binds to cell surface N-linked Neu5Ac, a derivative of sialic acid (SA) and common terminal end for complex glycans in mammalian cells^23^. Murine pancreatic cells are known to have high sialidase activity^24^, indicating the presence of Neu5Ac. AAV6 also displays affinity for heparin and may secondarily bind to HSPGs, but to a lesser degree than AAV2^21^. The addition of either heparin or neuraminidase (NM) should theoretically reduce AAV6 binding efficiencies as heparin could competitively bind to the virus while neuraminidase would cleave terminal Neu5Ac. Serotype 9 AAVs attach to cells via tethering to terminal galactose. In complex glycans, galactose comprises the layer below the SA cap and is generally inaccessible. Enzymatic cleavage of the cap, for example by NM, can remove neuraminic acids and expose galactose, potentially improving AAV9 binding efficacy^25^.

In this study, the infectivity of AAV2, AAV6 and AAV9 was tested on murine α- and β-cells. Transduction with AAV2 carrying the eGFP gene (thereafter GFP) of both cell types resulted in extensive expression whereas exposure to AAV6 or AAV9 resulted in limited GFP presence. Moreover, the interaction of β-cells with AAVs was modulated with heparin. Incubation of AAV2 with solubilized heparin before its addition to cells reduced transduction efficiency to near zero, pointing to the role of heparin as a competitive inhibitor in this process. NIH/3T3 murine fibroblast cells were also included in this study to investigate if our observations on pancreatic cells were applicable to non-pancreatic cells.

## MATERIALS AND METHODS

### Cell Culture

Pancreatic αTC1 clone 6 α-cells (αTC1-6, passage 16, ATCC, Manassas, VA), 293AD human embryonic kidney cells (HEK293, passage 42-61, Cell Biolabs, Inc., San Diego, CA), β-TC-6 mouse pancreatic β-cells (βTC; passage 40-69, ATCC), and NIH/3T3 (passage 33-40, ATCC) mouse fibroblasts were cultured at 37°C and 5% CO_2_ atmosphere. The αTC1-6 cells were maintained in Dulbecco’s Modified Eagle Medium (DMEM, Gibco, Grand Island, NY) containing 1 g/L glucose and supplemented with 10% fetal bovine serum (FBS, Corning, Corning, NY), 15 mM HEPES (Fisher BioReagents, Pittsburgh, PA), 0.1 mM MEM non-essential amino acids (Gibco), 0.02% w/v bovine serum albumin (fraction V cold-ethanol precipitated; Fisher BioReagents), and 100 U/mL penicillin and 100 µg/mL streptomycin (Cytiva, Marlborough, MA). All other cells were cultured in DMEM with 4.5 g/L glucose, 10% FBS, 100 U/mL penicillin and 100 µg/mL streptomycin. αTC1-6 cells were passaged every 5 days at a 1:3 splitting ratio. All other cells were passaged weekly by splitting the β-cells 1:3 and the NIH/3T3 cells 1:100. Media were exchanged every 2-3 days. Multi-well plates for culturing βTC and NIH/3T3 cells were coated with 1:50 Matrigel (Corning) and 1:500 human plasma fibronectin (MilliporeSigma, Burlington, MA) diluted in DMEM.

### Adeno associated virus production

For AAV production, HEK293 cells in T-225 flasks were transfected with the following plasmids (30 µg each): pHelper (Cell Biolabs), pAAV-GFP (Cell Biolabs), and a capsid plasmid. The capsid plasmids in this study are: pAAV 2/2 for AAV2 (Addgene #104963, Addgene, Cambridge, MA), pRepCap6 for AAV6 (Addgene #110770), or pAAV 2/9n for AAV9 (Addgene #112865). Calcium phosphate-based transfection was carried out^26^. Briefly, 2 M CaCl_2_ was added to a DNA mixture of pHelper, pAAV-GFP, and capsid plasmid (1:1:1 mass ratio). A 2x HEPES-buffered saline solution (50 mM HEPES, 1.5 mM Na_2_HPO_4_, 137 mM NaCl, pH 7.05) was then added slowly to the DNA/CaCl_2_ mixture followed by incubation at room temperature for 30 minutes. The resulting precipitate solution was added to HEK293 cells at 70% confluency in T-225 flasks. Media were exchanged after 24 hours. At 72 hours post-transfection, the cells were harvested and processed with an AAV purification kit (all serotypes; Takara Bio, San Jose, CA). The titer of the purified virus stocks was determined in viral genomes (vg)/ml using an AAV titration kit (Takara Bio).

### Cell transduction with AAV

The βTC cells were plated at 60,000 cells/well in coated 96-well plates. Experimental conditions tested were varied by the multiplicity of infection (MOI; vg/cell), heparin (Sigma-Aldrich, St. Louis, MO) and neuraminidase (NM; Sigma-Aldrich) treatment, as well as the AAV serotype (**Table 1**). Heparin was reconstituted in ultrapure water per the manufacturer’s instructions. Heparin and NM were diluted in Ca^2+^-/Mg^2+^-free PBS (Gibco).

**Table 1.**
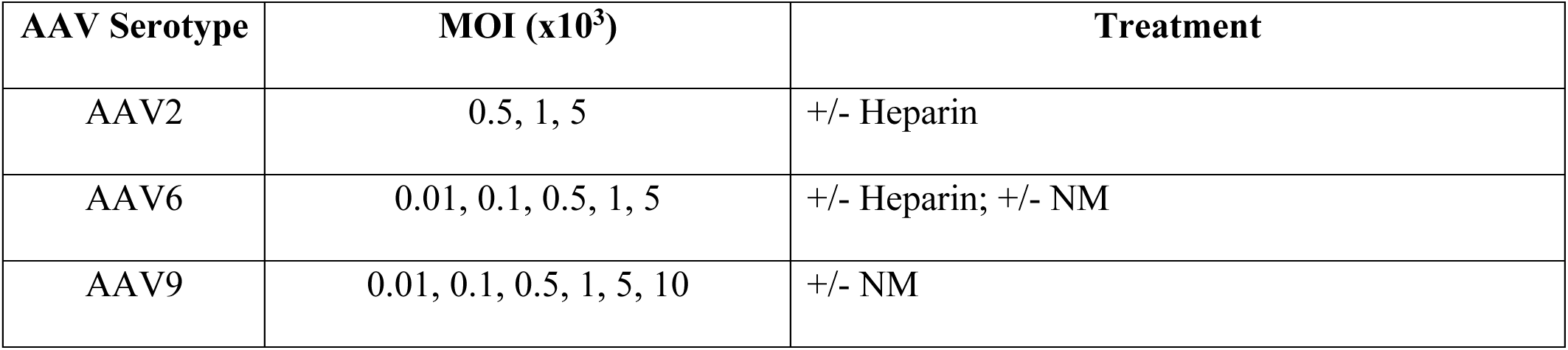
Summary of the experimental screening conditions reported in this work.

Heparin treatment entailed the incubation of the virus with 4.0×10^-7^ U heparin/vg at 37°C for 2 hours prior to cell infection. For exposure to NM, cells were incubated with 25 mU NM/mL diluted in PBS at 37°C for 2 hours before the addition of virus at a stated MOI. Media were exchanged 24 hours post-infection, and the cells were imaged daily for the next 5 days with an inverted microscope (Leica DM IL LED, Leica Microsystems, Buffalo Grove, IL) connected to a camera (Leica DFC3000G) and software (v.4.4 LASX, Leica).

### Flow Cytometry

The βTC and NIH/3T3 cells were plated on Matrigel-coated multi-well plates at a density of 2.6×10^5^ cells/cm^2^ and 1.7×10^5^ cells/cm^2^, respectively. Additionally, αTC1-6 cells were plated at 2.4×10^5^ cells/cm^2^. Five days post-transduction at MOIs ranging from 100 to 5,000, cells were washed using DMEM and treated with TrypLE Express Enzyme (Gibco). Following the addition of FBS-supplemented DMEM, the cells were subjected to centrifugation at 200x RCF for 5 minutes and resuspended in PBS prior to flow cytometry analysis (Attune NxT, ThermoFisher Scientific, Waltham, MA). The data were analyzed using the FCS Express software (v. 7, De Novo Software, Pasadena, CA). Cells were designated as positive for GFP expression if their emitted fluorescence was >99% of the corresponding control cultures (virus-free cells). A fraction of the resuspended cells was counted and their viability was assessed using a BioRad TC20 Automated Cell Counter (BioRad, Hercules, CA) after incubation with Trypan Blue (Sigma-Aldrich).

### Statistical Analysis

Flow data are expressed as mean ± standard deviation (SD) from independent samples analyzed in quadruplicates. ANOVA and the *posthoc* Tukey test were performed using Prism (v. 10.2.3, GraphPad Software, La Jolla, CA). Values of p<0.05 were considered as significant.

## RESULTS

### AAV2 exhibits tropism for pancreatic beta-cells that is inhibited by soluble heparin

The heparan sulfate proteoglycan (HSPG) cell surface receptor is a common binding site for different virus types. Specific to AAVs, AAV2 utilizes HSPG as its primary partner for initial interactions with target cells. Heparin, which is found in the cytoplasmic granules of mast cells, exhibits a highly similar molecular structure to heparan sulfate, which is ubiquitously displayed on the surface and extracellular matrix of all cells^27^. Both are composed of the same monomeric building blocks but in different proportions of the various components^28^. Previous studies have shown that heparin-based chromatography retains AAV2 as well as AAV6^21^, yet only AAV2 shows competitive interaction with heparin in airway epithelial cells. We hypothesized that AAV2 pre-treated with heparin would have its binding sites occupied, leading to reduction of the transduction efficiency of pancreatic β-cells.

Prior to our screening study, we conducted a toxicity screen to ensure that heparin does not induce any negative side effects in βTC cells. Cells were exposed to media containing 2400 U heparin/mL for 24 h, the maximum possible heparin concentration corresponding with an infection of 10,000 vg/cell. No changes in cell morphology, viability, or growth rate were noted compared to the control, indicating no inherent cytotoxicity or other adverse effects due to heparin exposure (data not shown).

To test our hypothesis, βTC β-cells were exposed to AAV2 at varying multiplicities of infection (MOI; vg/cell) with and without heparin pre-treatment of the viral particles. The βTC cells mirror many attributes of native β-cells including their GSIS response and ability to self-assemble into islet-like structures^29^. Cells were exposed to 500 – 5,000 vg/cell with or without pre-incubation of the virus with heparin. Five days post-infection, cells at all MOI showed similar morphology and confluency close to 100%, indicating negligible virus-induced toxicity (**Fig. 1A-C**). At 1,000 vg/cell (**Fig. 1B**), a low level of GFP expression was observed irrespective of heparin treatment, suggesting that this MOI was unable to induce sufficient transduction for appreciable expression. However, noticeable GFP expression was seen at 5,000 vg/cell without incubation with heparin (**Fig. 1C**). Cells receiving AAV2 pre-treated with heparin exhibited less GFP expression than cells exposed to AAV2 alone. This finding points to the competitive action of heparin when binding to AAV2 resulting in reduced transduction of the β-cells.

**Figure 1.**
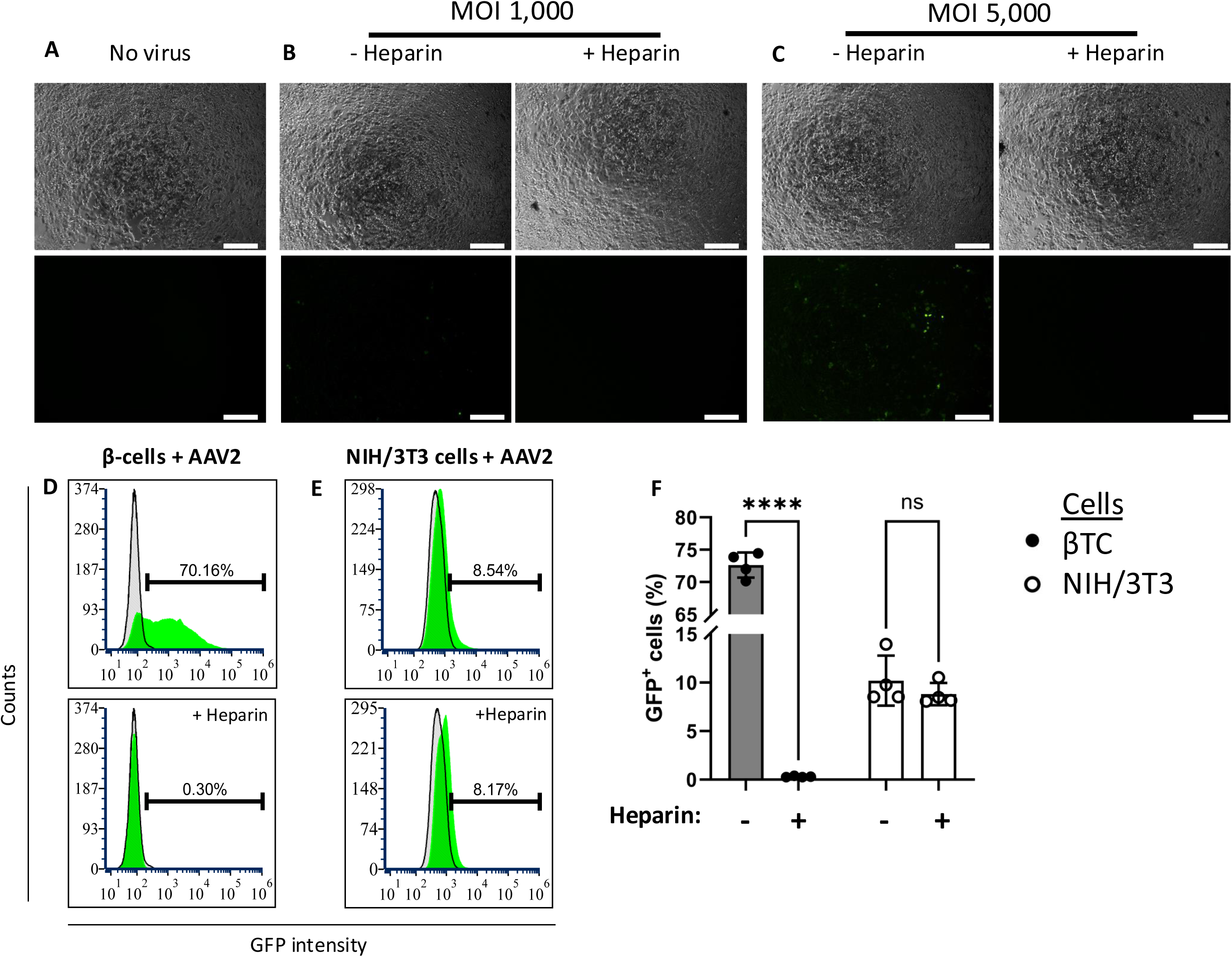
Cell transduction with AAV2. (A) βTC cell control cultures or transduced with (B) 1,000 or (C) 5,000 vg/cell of AAV2 incubated without (-Heparin) or with heparin (+Heparin). Top row: Brightfield images. Bottom row: Fluorescence images (GFP). Images were taken at 120 hours post-infection. Bars: 200 μm. (D-E) Representative flow cytometry results showing the fraction of GFP^+^ cells (green curve) upon transduction of (D) βTC cells and (E) NIH/3T3 cells, with AAV2 at 5,000 vg/cell without heparin (top graphs) or with heparin (bottom graphs). Gray curves represent untransduced cells. (F) Summary of flow cytometry results of βTC and NIH/3T3 cells infected with AAV2 at 5,000 vg/cell. ****: p<10^-4^, ns: not significant.

Given that βTC cells showed GFP fluorescence after infection with AAV2 carrying the GFP transgene, the fraction of GFP-positive cells was quantified by flow cytometry to determine the efficiency of transduction. The same samples were also counted and assessed for viability to check virus-induced cytotoxicity. The βTC cells were transduced at 5,000 vg/cell with and without heparin pre-treatment of the virus. In parallel, NIH/3T3 murine fibroblasts were identically transduced as a control to examine if AAV2 infects systemically or has a specific tropism towards pancreatic cells. Moreover, αTC α-cells were transduced at 500 and 5,000 vg/cell without heparin to confirm if AAV2’s tropism extends to other pancreatic cells (**Fig. 2**).

**Figure 2.**
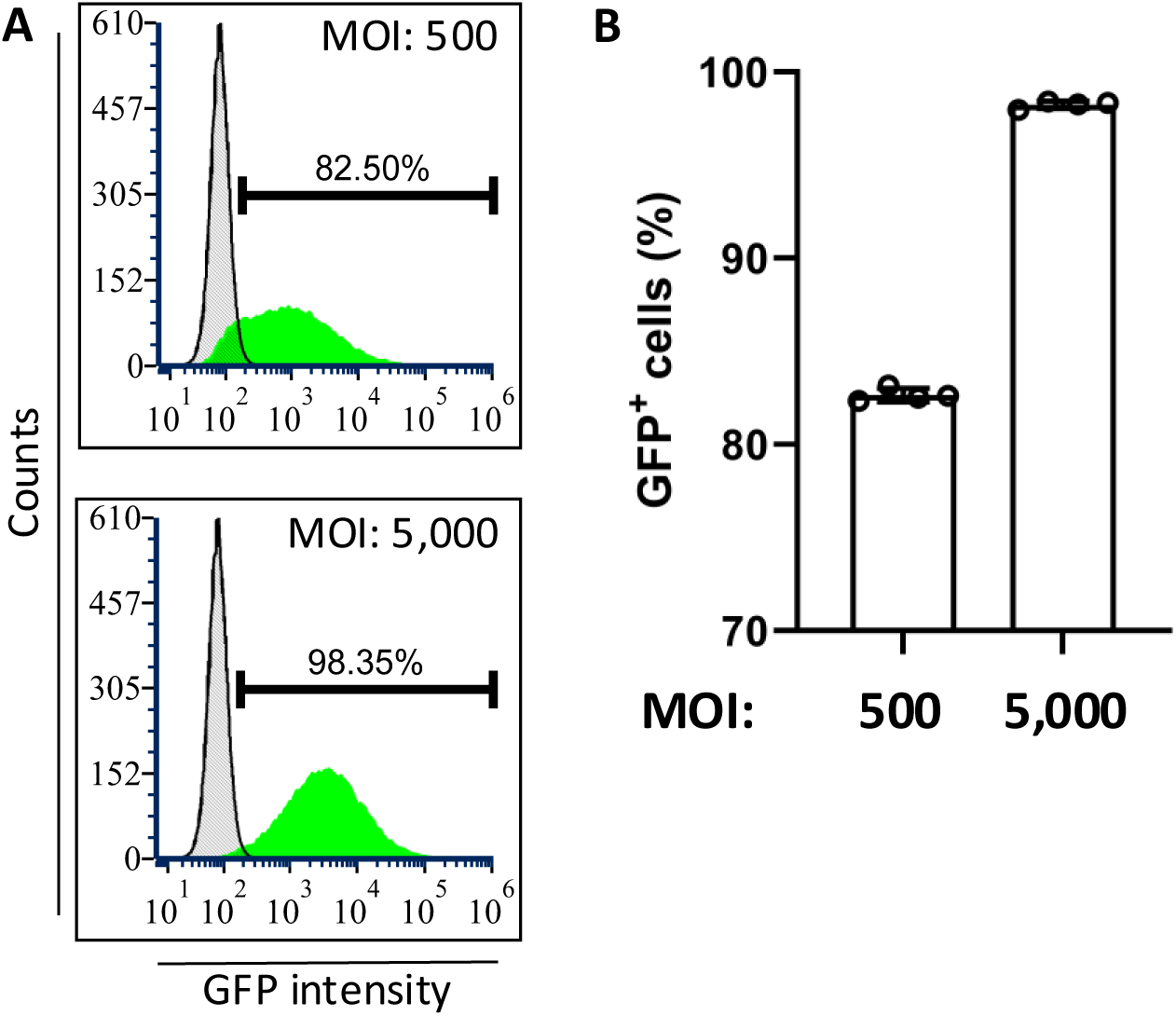
Flow results of AAV2 transduction in αTC1-6 cells. (A) Representative flow cytometry results of αTC cells infected with 500 or 5,000 vg/cell of AAV2. Curves correspond to infected (green) and untransduced (control; gray) cells. (B) Summary of flow cytometry results for αTC cell transduction with AAV2. Results are shown as mean ± SD.

On average 72.6% ± 1.9% of AAV2 transduced βTC cells were positive for GFP expression (**Fig. 1D**). Pre-treatment of the virus with heparin led to complete inhibition of productive infection as indicated by the lack of GFP expression, resulting in a significant difference between the two groups (p < 10^-4^, n=4). In contrast, NIH/3T3 fibroblasts infected with the AAV2 at 5,000 vg/cell displayed an average of 10.2% ± 2.6% and 8.2% ± 1.2% GFP^+^ cells without and with heparin, respectively (**Figs. 1E-F**; p=0.674, n=4). AAV2 transduction of α-cells proved to be more efficient, with 83.2% ± 0.4% and 98.2% ± 0.2% of infected cells displaying GFP at 500 and 5,000 vg/cell, respectively (**Fig. 2**). The viability of βTC and NIH/3T3 cells exposed to AAV2 at 5,000 vg/cell ranged between 75-95% and appeared to be the lowest (70 %) for NIH/3T3 cells infected with heparin-treated AAV2 (**Fig. 3A**). The αTC1-6 cells showed no appreciable changed in viability with exposure to AAV2 (**Fig. 3B**).

**Figure 3.**
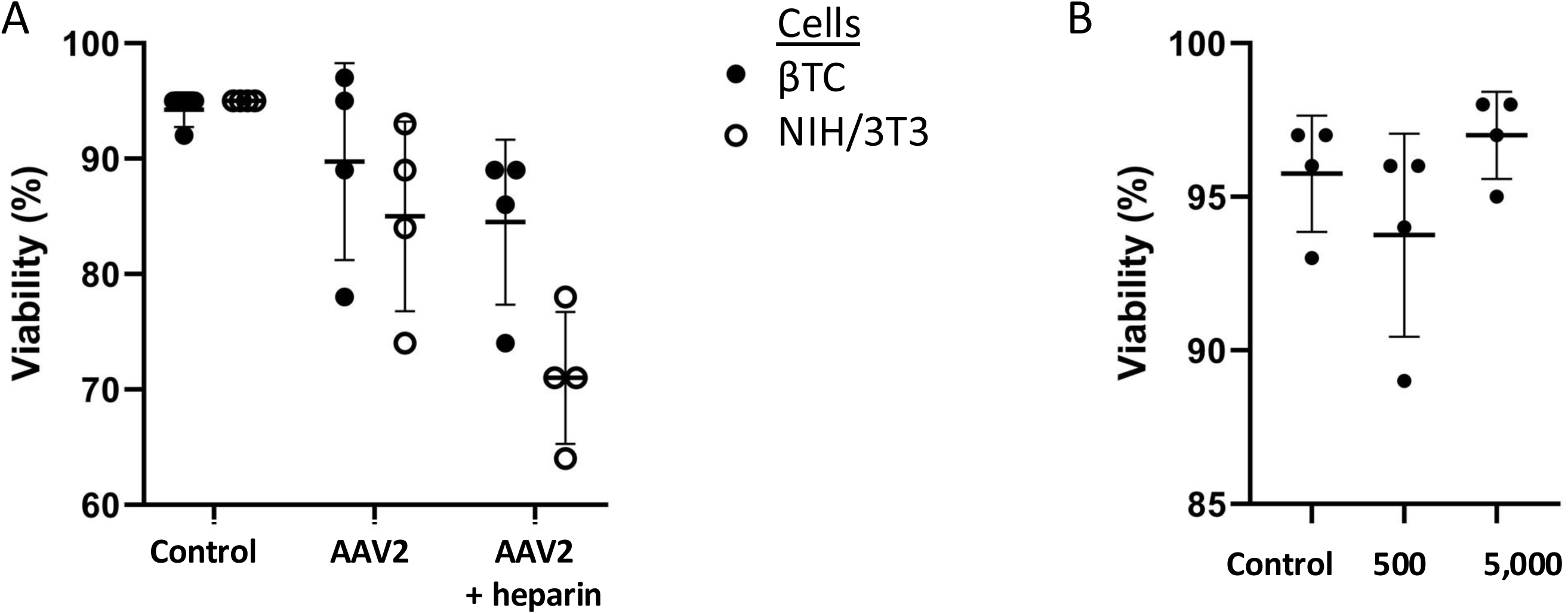
Viability results of AAV2 transduction in (A) βTC and NIH/3T3 cells (5,000 vg/cell), and (B) αTC1-6 cells at MOI as indicated. Results are shown as mean ± SD.

### AAV6 does not induce efficient infection of β-cells

AAV6 utilizes N-linked SA moieties as its primary binding receptor and HSPG as a co-receptor. Hence, in addition to virus pre-incubation with heparin, transduction of β-cells with AAV6 was tested after cell treatment with the glycoside hydrolase neuraminidase (NM) which cleaves N-linked SAs. Both variables were investigated simultaneously to assess their effect on AAV6 infection of βTCs. Between 500 – 5,000 AAV6 vg/cell were added to cultured βTCs (**Fig. 4, Table S2**). All conditions were tested in triplicate.

**Figure 4.**
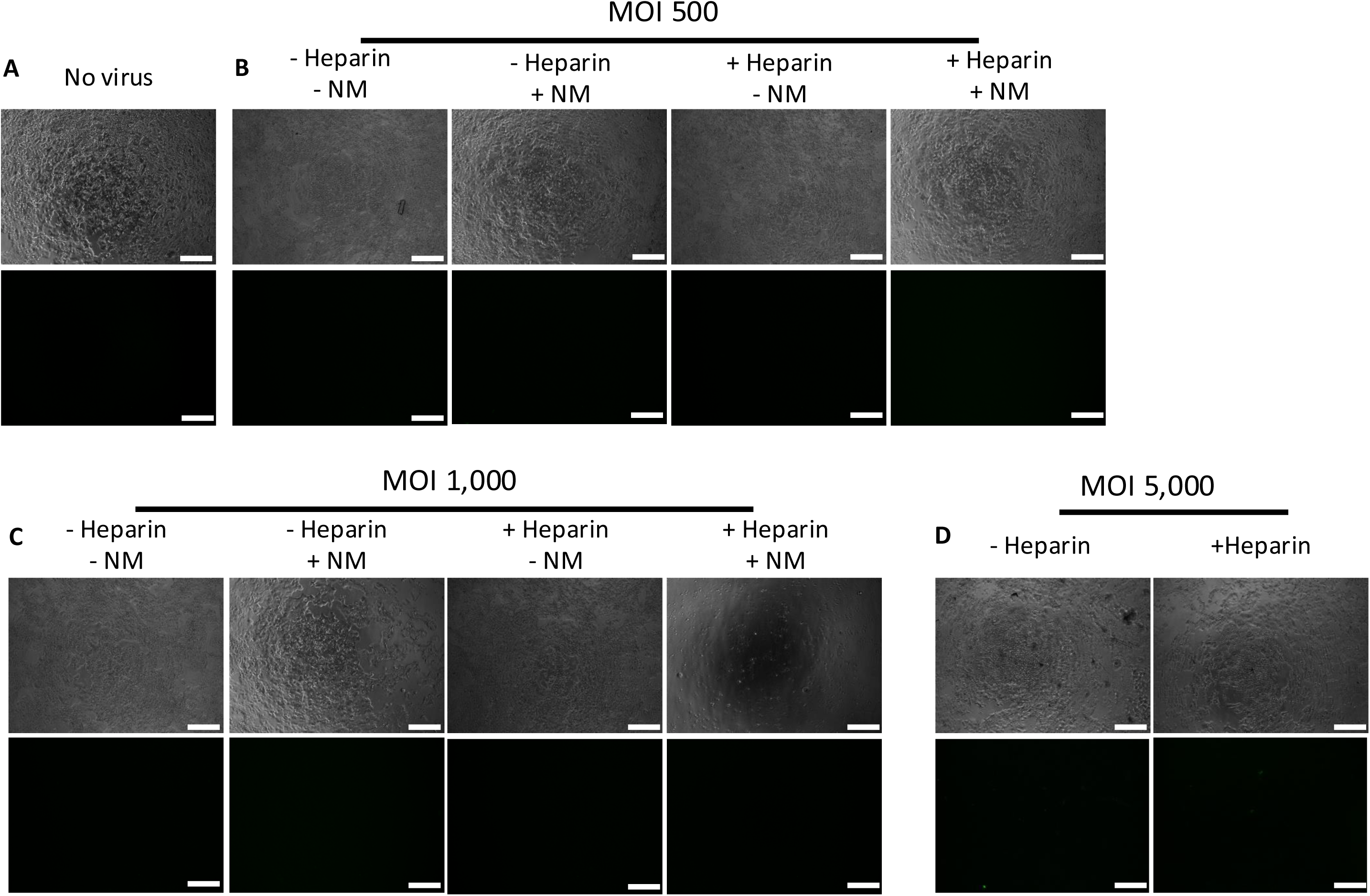
AAV6 transduction of β-cells. Images of cultured βTC cells (A) without virus (control), or with AAV6 at (B) 500, (C) 1,000, and (D) 5,000 vg/cell. Treatments with NM and heparin are indicated. Top rows: Brightfield images. Bottom rows: Fluorescence images (GFP). Images were taken at 120 hours post-infection. Bars: 200 μm.

Prior to viral testing, a toxicity screen of NM was conducted on both βTC and NIH/3T3 cells. Cells were exposed to 25 mU NM/mL PBS for 2 hours, followed by media exchanging into fresh complete medium and recovering for 22 hours. After the initial 2-hour exposure, βTC cell morphology was significantly altered with cells assuming a round shape with partial loss of adhesion. After the recovery period, all cells had spread out again and appeared equally healthy to the control. This recoverable morphology change indicates that SA cleavage has an effect on βTC cell adhesion. The NIH/3T3 cells did not display this same effect (**Fig. S1**).

For βTC cells exposed to AAV6 alone at 1,000 vg/cell or less, they grew to confluency but no GFP expression was observed (**Figs. 4A-C**). When infected with 5,000 vg/cell, β-cells visibly expressed GFP but cell loss was noted indicating toxicity (**Fig. 4D**). Cell treatment with NM followed by infection with AAV6 at all tested MOIs resulted in no GFP expression (**Fig. 4**). Incubation with NM followed by AAV6 transduction at 1,000 vg/cell resulted in cultures with lower confluency compared to those with NM treatment and MOI of 500 vg/cell (**Figs. 4C-B**), suggesting that incubation with a higher viral load leads to greater cytotoxicity or cell loss when co-treated with NM. This is additionally supported by the lack of viable cells in wells treated with NM and infected at 5,000 vg/cell of AAV6 (**Fig. S2**).

When βTC cells were combined with heparin-treated AAV6 at lower MOI, they displayed no morphological changes, cytotoxicity, nor GFP fluorescence. βTC cells with 5,000 vg/cell of AAV6 combined with heparin showed GFP expression, albeit at a reduced level compared to virus alone along with a higher level of cytotoxicity (**Fig. 4D**). However, the combination of AAV6 at MOI of 1,000 (**Fig. 4C**) and 5,000 (**Fig. S2**) pre-treated with heparin and NM-treated cells resulted in widespread death. Using the same treatment at an MOI of 500 showed no overt negative effect on cell viability (**Fig. 4B**). These results indicate the AAV6’s inability to meaningfully induce GFP expression without an accompanied loss of viability in β-cells.

### Infection with AAV9 results in differential transgene expression in β- and α-cells

The AAV9 serotype tethers to the terminal galactose of cell surface glycans to initiate infection. Enzymatic cleavage of the glycan cap above the galactose, for instance by NM, may increase exposure of galactose and potentially enhance transduction. Published studies confirm as much, with NM treatment of neonatal rat ventricular muscle cells increasing their transduction with AAV9^25^. Thus, similar to the aforementioned experiments with AAV6, NM pretreatment of murine βTC cells was tested to see its effect on AAV9 transduction efficiency.

Infections were carried out at a MOI range of 50-10,000 vg/cell (**Fig. 5, Table S3**). Five days after AAV9 addition, all virus-only cultures showed similar cell morphology and confluency at close to 100% as cultures without infection, indicating minimal cytotoxicity due to the presence of virus. Transductions at 1,000 vg/cell and above without NM displayed visible GFP expression (**Figs. 5C-E**). For β-cells treated with NM, cell coverage decreased significantly with increasing AAV9 MOI, indicating a loss of adhesion and/or induced cytotoxicity that scaled with added virus. From 100 to 1,000 vg/cell, GFP expression was visibly elevated upon treatment with NM compared to virus only-treated cells. At 5,000 vg/cell and above, no cells were attached to the wells after incubation with NM (not shown). Based on these results, MOI of 100 and 500 vg/cell were selected to further assess quantitatively the transduction efficiency of β-cells using flow cytometry. Cells were also tested at 5,000 vg/cell of AAV9 without NM for comparison.

**Figure 5.**
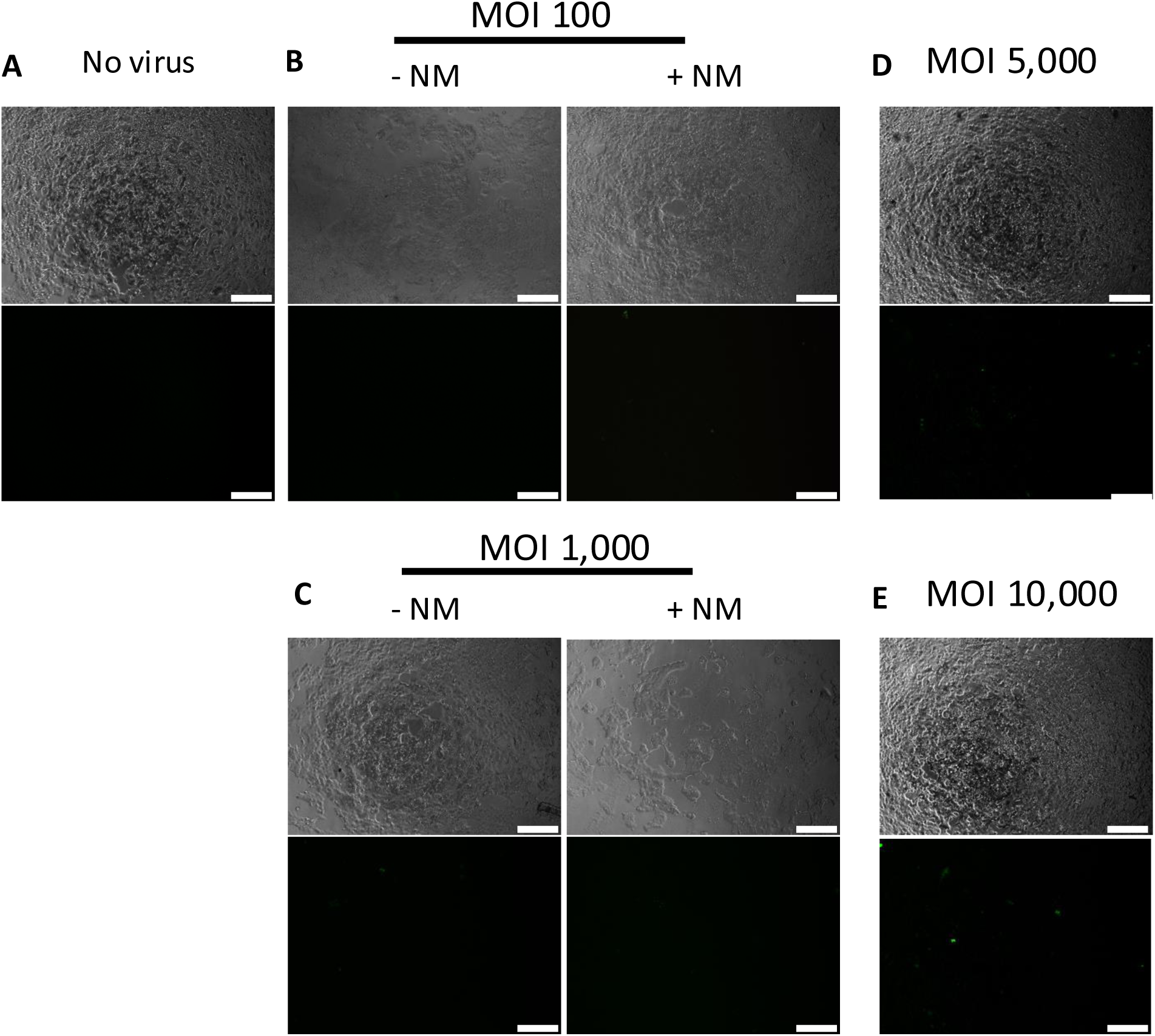
AAV9 transduction of β-cells. Images of cultured βTC cells (A) without virus (control), or with AAV9 at (B) 100, (C) 1,000, (D) 5,000, and (E) 10,000 vg/cell. Treatment with NM is indicated. Top rows: Brightfield images. Bottom rows: Fluorescence images (GFP). Images were taken at 120 hours post-infection. Bars: 200 μm.

For β-cells transduced with AAV9 at 100 vg/cell, the average fractions of GFP^+^ cells were 2.3% ± 0.2% and 1.5% ± 0.1% with and without prior NM incubation, respectively (**Figs. 6A-B, 6K**). At 500 vg/cell with and without NM treatment the corresponding fractions were 2.4% ± 0.4% and 1.8 % ± 0.4%, respectively (**Figs. 6C-D, 6K**). In both cases, no significant difference was found between the AAV9 only and combined virus/NM conditions (MOI 100: p = 0.186, n=4; MOI 500: p = 0.525, n=4). Increasing the MOI to 5,000 vg/cell significantly improved GFP expression, with 8.2% ± 0.8% being GFP^+^ (against all other conditions p < 10^-4^, n=4) (**Fig. 6E**). No differences in viability were found across all conditions (**Fig. 7A**).

**Figure 6.**
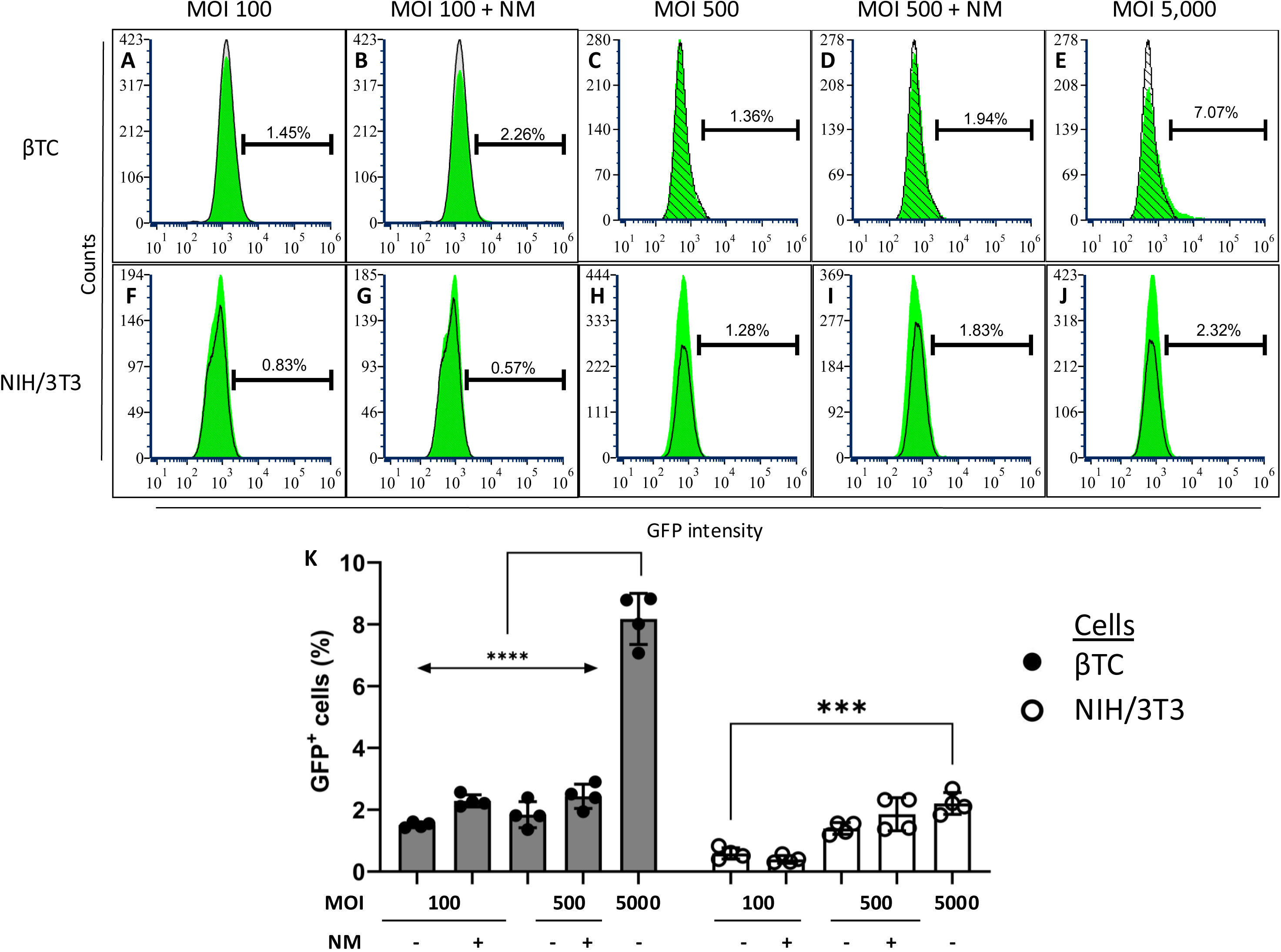
Representative results for AAV9-infected (A-E) βTC cells and (F-J) NIH/3T3 cells. The MOI and NM treatment are indicated. Curves correspond to untransduced (gray) and infected (green) cells. (K) Summary of flow cytometry results of βTC and NIH/3T3 cells infected with AAV9. Results are shown as mean ± SD.***: p<10^-3^, ****: p<10^-4^, ns: not significant.

**Figure 7.**
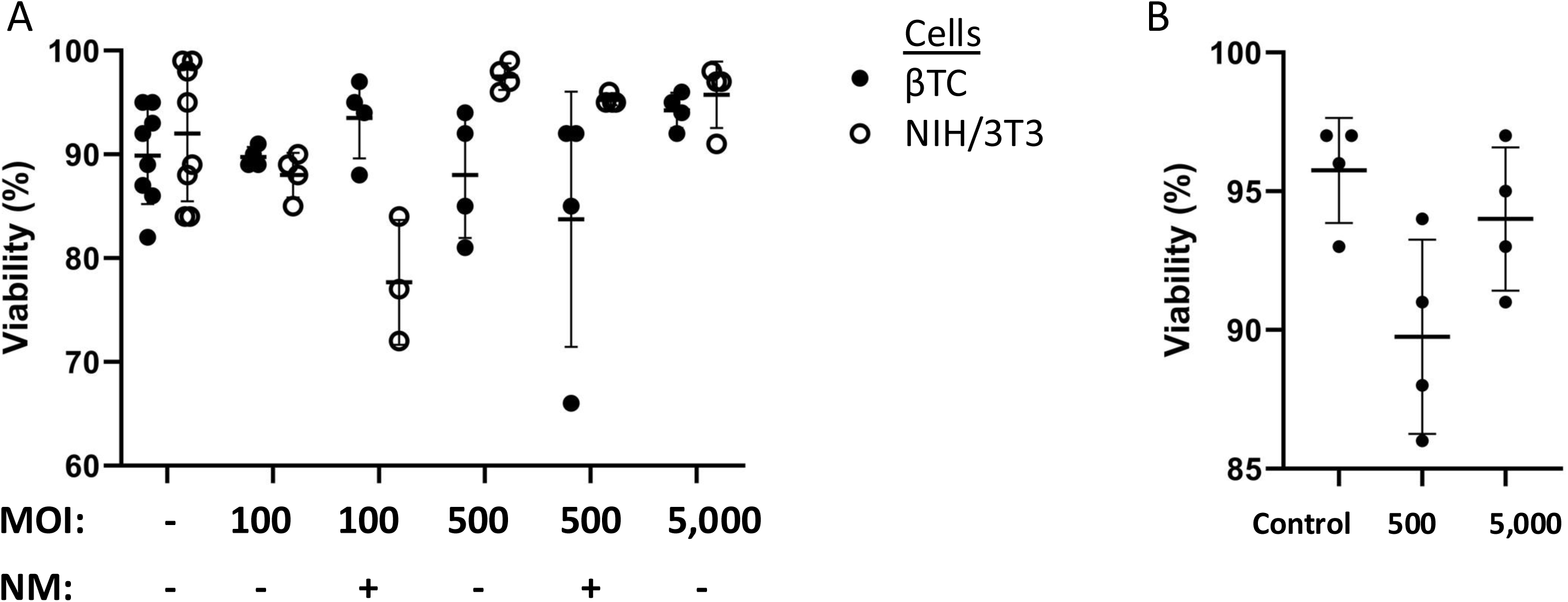
Viability of (A) βTC and NIH/3T3 cells, and (B) αTC1-6 cells, upon transduction with AAV9 at MOI as indicated. Control represents untransduced cells. Results are shown as mean ± SD.

As with the quantitative analysis of AAV2 infection, NIH/3T3 and αTC1-6 cells were also transduced with AAV9 here and examined by flow cytometry. NIH/3T3 cells displayed comparable levels of GFP^+^ cells to the β-cells. At 100 vg/cell, 0.6% ± 0.2% and 0.4% ± 0.1% of cells were GFP^+^, without and with NM treatment respectively (**Figs. 6F-G**). Similarly, at 500 vg/cell, the corresponding fractions of GFP^+^ cells were 1.4% ± 0.2% and 1.9% ± 0.5%, respectively (**Figs. 6H-I**). Again, no significant difference was found between the AAV9 only and combined AAV9/NM conditions (MOI 100 vg/cell: p=0.999, n=4; MOI 500 vg/cell: p=0.812, n=4). Escalating the MOI to 5,000 vg/cell slightly increased the percentage of GFP-expressing cells (2.2% ± 0.4%; **Fig. 6J**).

Interestingly, αTC1-6 cells showed the highest affinity for AAV9 of the cell types tested. At 500 vg/cell, 8.1% ± 0.4% of cells were GFP^+^ and 38.8% ± 1.2% at 5,000 vg/cell (**Fig. 8**). No significant differences in viability were detected compared to the control samples (**Fig. 7B**).

**Figure 8.**
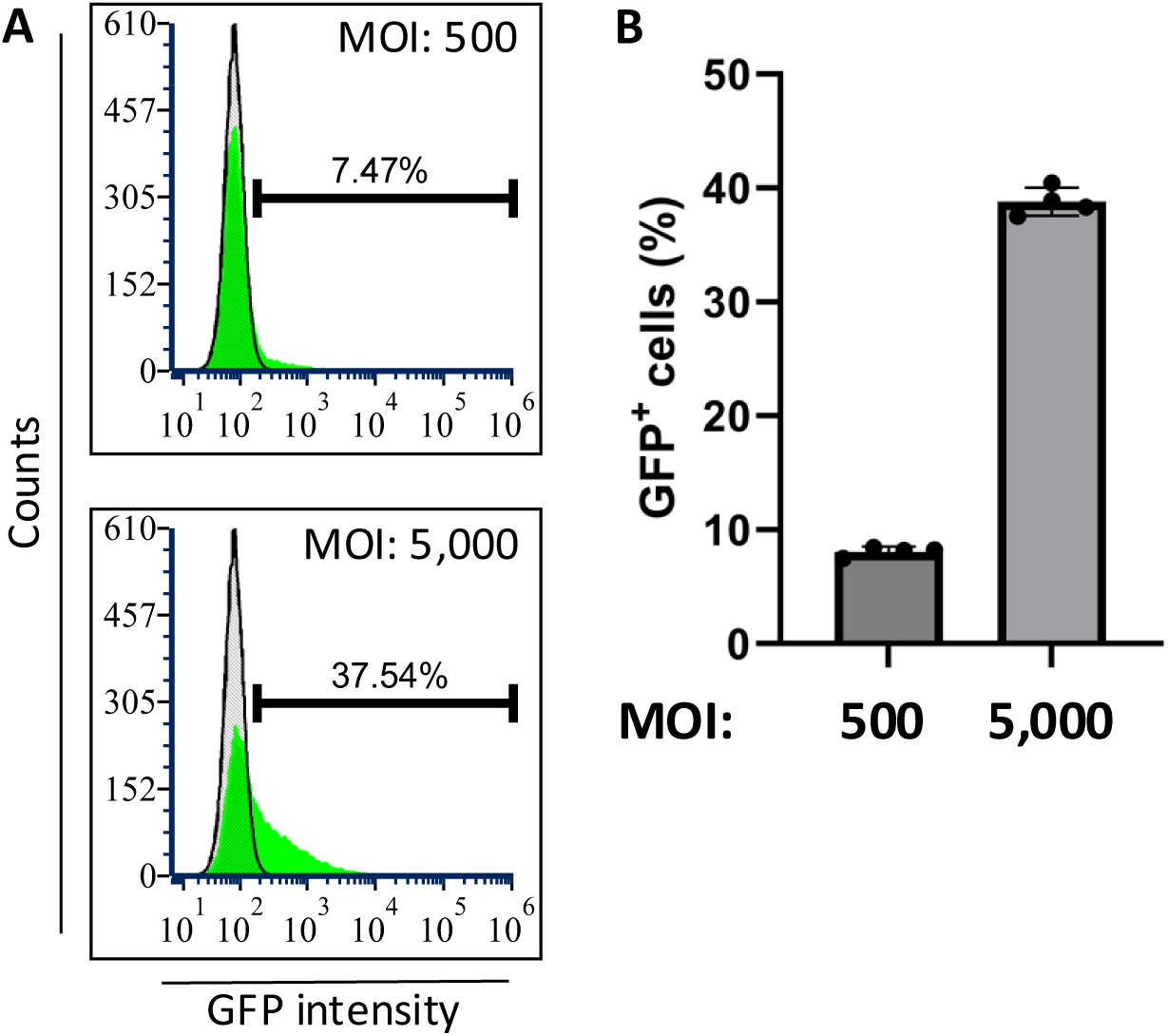
Flow results of AAV9 transduction in αTC1-6 cells. (A) Representative flow cytometry results of αTC cells infected with 500 or 5,000 vg/cell of AAV9. (B) Summary of flow cytometry results for αTC cell transduction with AAV9. Results are shown as mean ± SD.

## DISCUSSION

AAVs have gained attention as a powerful tool for cell and gene therapies. Despite their limited packaging capacity compared to alternative gene delivery vehicles such as lentiviruses, AAVs excel due to their robust safety^9^ and superior manufacturability^30^. AAV8 in particular has shown high tropism for pancreatic cell types, and in various studies it has been applied as a genetic vector in the context of diabetes^10–13^. With AAV8 being the only major serotype commonly used with pancreatic endocrine cells however, we endeavored here to expand the repertoire of available serotypes by testing AAV2, AAV6, and AAV9 on these cells.

Prior to testing the AAVs, we conducted studies for both heparin (not shown) and NM (**Fig. S1**) to assess potential cytotoxic effects and ensure that our treatment results were due only to either viral exposure or combinatorial effects. When β-cells were exposed to heparin for 24 hours, they showed no visible morphological changes or death. Treatment with NM for 2 hours however resulted in the cells rounding but they fully recovered, reverting to their normal morphology within a day after NM removal. No loss of cells was observed and the overall plate confluency was comparable to the initial coverage prior to NM exposure. These morphological changes indicate that the cells may be prone to loss of adhesion with NM-mediated cleavage of SA on their surface. When the NIH/3T3 cells were tested with heparin and NM, no toxicity or morphological changes were noted (**Fig. S1**), indicating different surface glycosylation patterns from β-cells.

Our results revealed that AAV2 has a natural affinity for β-cells, and the preservation of culture confluency suggests minimally induced cytotoxicity. The extensive expression of GFP points to the presence of HSPGs as moieties of AAV2 interaction on the surface of β-cells. This is further corroborated by the lack of GFP expression in cells exposed to heparin-treated AAV2, indicative of competitive inhibition. The flow cytometry analysis confirmed that AAV2 transduction was efficient with ∼72% of the β-cells expressing GFP. In contrast, cells exposed to AAV2 incubated with heparin showed no significant differences in GFP fluorescence compared to the untransduced control cells. AAV2 affinity was additionally confirmed to extend to pancreatic αTC1-6 α-cells, with nearly 98% of the infected cells expressing GFP at 5,000 vg/cell. As of this time, we have not tested αTC1-6 cells with heparin-incubated AAV2 and so cannot conclusively state that AAV2 interacts with α-cells through HSPGs, but given the observed high rate of infection we believe that this is still the most likely mode of interaction.

The tropism of AAV2 does not extend to the tested murine fibroblast cells. Approximately 8-10% of transduced NIH/3T3 cells were GFP^+^, irrespective of pre-treatment of the virus with heparin. We therefore assume that AAV2 does not infect this fibroblast line via HSPGs but instead through some unknown secondary binding site. Alternative candidates include fibroblast growth factor receptor 1 (FGFR1), αVβ5 and α5β1 integrins, hepatocyte growth factor receptor (HGFR), laminin receptor (LR), and CD9^31^. The presumption is that pancreatic cell types have similar HSPG glycosylation patterns to one another which enhances AAV2 transduction efficiency, but the glycosylation profile does not carry over to other cell types. This positively suggests that an AAV2 infection *in vivo* would not be systemic and instead have some tropism towards pancreatic islets, presuming that the glycosylation profiles remain similar *in vitro* and *in vivo*. An interesting follow-up study would be to examine if AAV2 exhibits affinity for non-islet pancreatic tissue, such as acinar and ductal cells.

Unlike AAV2, AAV6 used to infect β-cells resulted in GFP expression at 5,000 vg/cell, but there was loss of confluency and a change in cell morphology. Infected βTC cells were visibly smaller, indicating that AAV6 induces a degree of cytotoxicity. Despite the genetic similarity to AAV2^31^, AAV6 could be toxic through various pathways including triggering cell apoptosis^32^.

Cell death due to AAV6 was further exacerbated by treatment with NM prior to initiating transduction. While the morphology of cells was unaffected, increasing the MOI resulted in lower overall culture confluency. This dose-dependent loss of cells could be occurring because the virus occupies cell-surface binding sites normally reserved for adhesion to the substratum. N-linked glycans have been shown to influence cell adhesion and migration, with truncated glycans missing SA caps having reduced migratory abilities^33^. While NM exposure alone would temporarily inhibit cell adhesion, βTCs showed that they re-adhere fully after NM removal. It is possible that NM treatment immediately followed with viral infection could result in all adhesive binding sites becoming either occupied or cleaved, resulting in cell detachment. Regardless, GFP expression in NM-treated wells was not visually detected at any MOI. This confirms that NM does not enhance AAV6 transfection efficiencies, but we were unable to prove whether NM actively suppresses transduction. Even for cells infected with heparin-treated AAV6, quantification of the GFP expression by flow cytometry was not possible due to the low cell viability.

It should be noted that AAV6 has an affinity for the Neu5Ac variant of SA. The presence of the Neu_5_Gc variant in murine cell lines^34^ may be reducing the number of possible binding sites. Increased transduction efficiencies may be possible with human pancreatic cells as humans cannot produce Neu_5_Gc^35^, so all complex glycans will be capped with Neu5Ac.

Like AAV2 and AAV6, β-cells treated with AAV9 at 5,000 vg/cell and above displayed GFP expression. At lower MOI, GFP fluorescence was visibly enhanced when cells were pre-treated with NM prior to initiating viral infections. At MOI of 100 vg/cell, the NM enhancement was especially visible, with no GFP expression in virus-only treated cells compared to GFP^+^ cells in NM/virus-treated wells. This is aligned well with published literature and our hypothesis that NM cleavage of SA caps enhances transduction efficiencies by exposing the terminal galactose to which AAV9 binds.

As with AAV6 however, treating β-cells with NM resulted in the same virus dose-dependent loss of cells. Given this loss, flow cytometry analysis was carried out at MOI of 100, 500, and 5,000 vg/cell. For both β-cells and fibroblasts, the number of GFP^+^ cells tends to increase with the MOI in NM-treated cultures compared to those exposed to the virus only. However, this difference is both statistically and practically insignificant. The only major jumps in the percentage of GFP^+^ cells were noted when the MOI increased to 5,000 vg/cell for both the βTC and αTC1-6 cells. The natural tropism for pancreatic islet cells is lower for AAV9 compared to AAV2, achieving only ∼8% and 39% GFP^+^ cells for β- and α-cells, respectively. Further augmentation in the fraction of GFP^+^ cells is certainly possible by boosting the MOI, however it would still require 2- to 5-fold more virus compared to using AAV2. The percentage of GFP^+^ NIH/3T3 cells was ∼2% at 5,000 vg/cell, indicating that AAV9 treatment *in vivo* would not be systemic. These differences in infectivity may be attributed to variations in glycosylation profiles, with cells potentially displaying simpler glycans without SA caps having stronger binding with AAV9. Glycosylation profiles of these cell lines would need to be studied for confirmation.

It should be mentioned that only a single line of β-cells was employed in this work. While βTC cells are representative of native murine β-cells, and most likely the overall patterns of infectivity hold true across different lines, small differences in glycosylation patterns among alternative lines cannot be ruled out.

Overall, our results show that AAV2 is a promising tool for delivering transgenes to pancreatic β- and α- cells, with a 72% and 98% transduction efficiency respectively. In contrast, the AAV serotypes 6 and 9 displayed poor infectivity of the cells. By introducing competitive inhibition through solubilized heparin, we have additionally shown that AAV2 primarily targets HSPGs expressed on the surface of β-cells. The use of this serotype should help expand the repertoire of transduction options beyond AAV8 and aid in optimizing future gene and cell therapy treatments for diabetes.

## ACKNOWLEDGEMENTS

Funding support has been provided by the National Science Foundation (NSF; CBET-1951104, CBET-2015849, CBET-2326510) to EST.

## AUTHOR CONTRIBUTIONS

S. Jeyabalan and V. Ahuja performed experiments, collected and analyzed data, and contributed to writing the manuscript; E.S. Tzanakakis analyzed data, acquired funding, and contributed to the writing and review of the manuscript.

## LIST OF ABBREVIATIONS

AAV: Adeno-associated virus
DMEM: Dulbecco’s modified Eagle medium
eGFP: Enhanced green fluorescent protein
FBS: Fetal bovine serum
HEPES: 4-(2-Hydroxyethyl)piperazine-1-ethanesulfonic acid
MOI: Multiplicity of infection
Neu5Ac: N-Acetylneuraminic acid
NM: Neuraminidase
PBS: Phosphate buffered saline
SA: Sialic Acid
VG: Viral genome

## FIGURE LEGENDS

**Figure S1.**
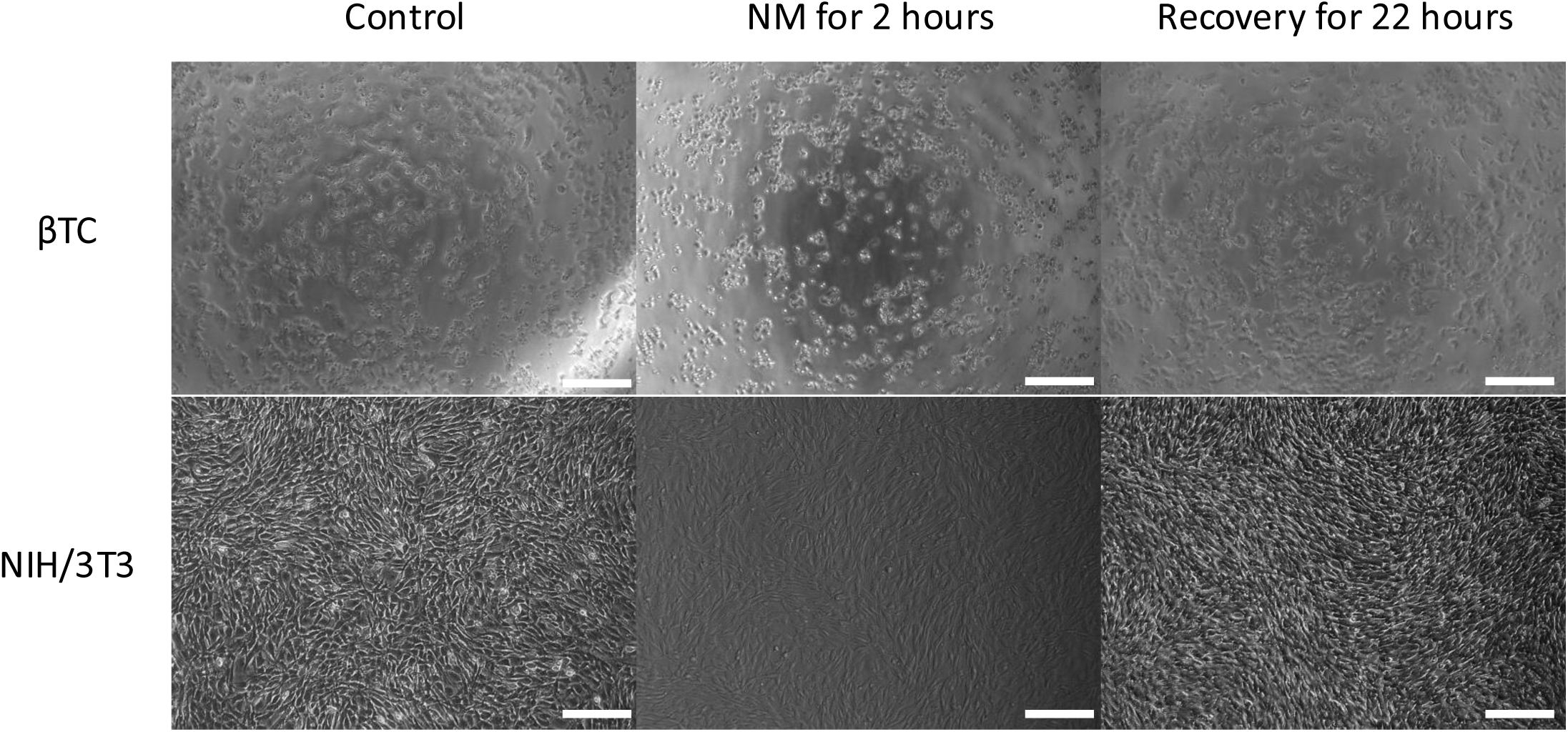
Brightfield images of (A) βTC and (B) NIH/3T3 cells treated with NM. Images show cells prior to NM exposure (left), 2 hours post-exposure (middle), and 22 hours after the recovery period in complete medium (right).

**Figure S2.**
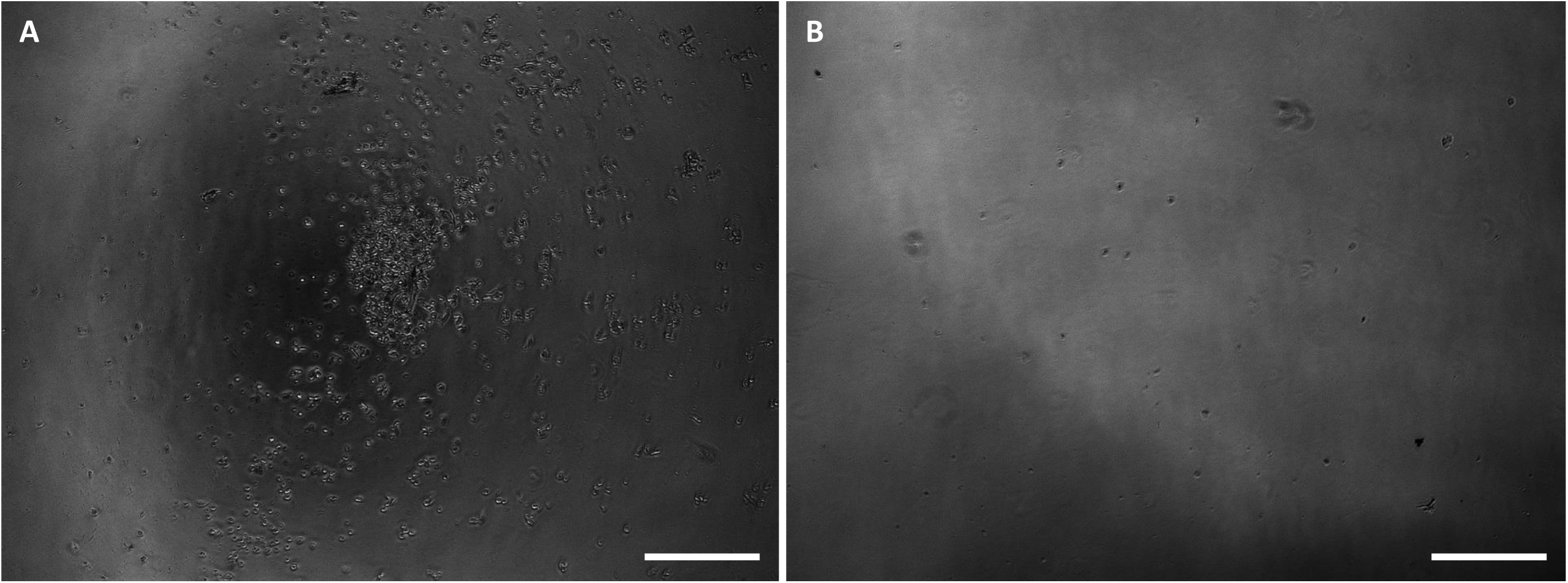
Brightfield images of AAV6 transduction of βTC cells at 5,000 vg/cell. (A) βTC cells pre-treated with NM prior to viral infection. (B) βTC cells pre-treated with NM prior to transduction with heparin-treated virus. Images were taken 48 hours post-infection. Bars: 200 μm.

## SUPPLEMENT

**Table S1.**
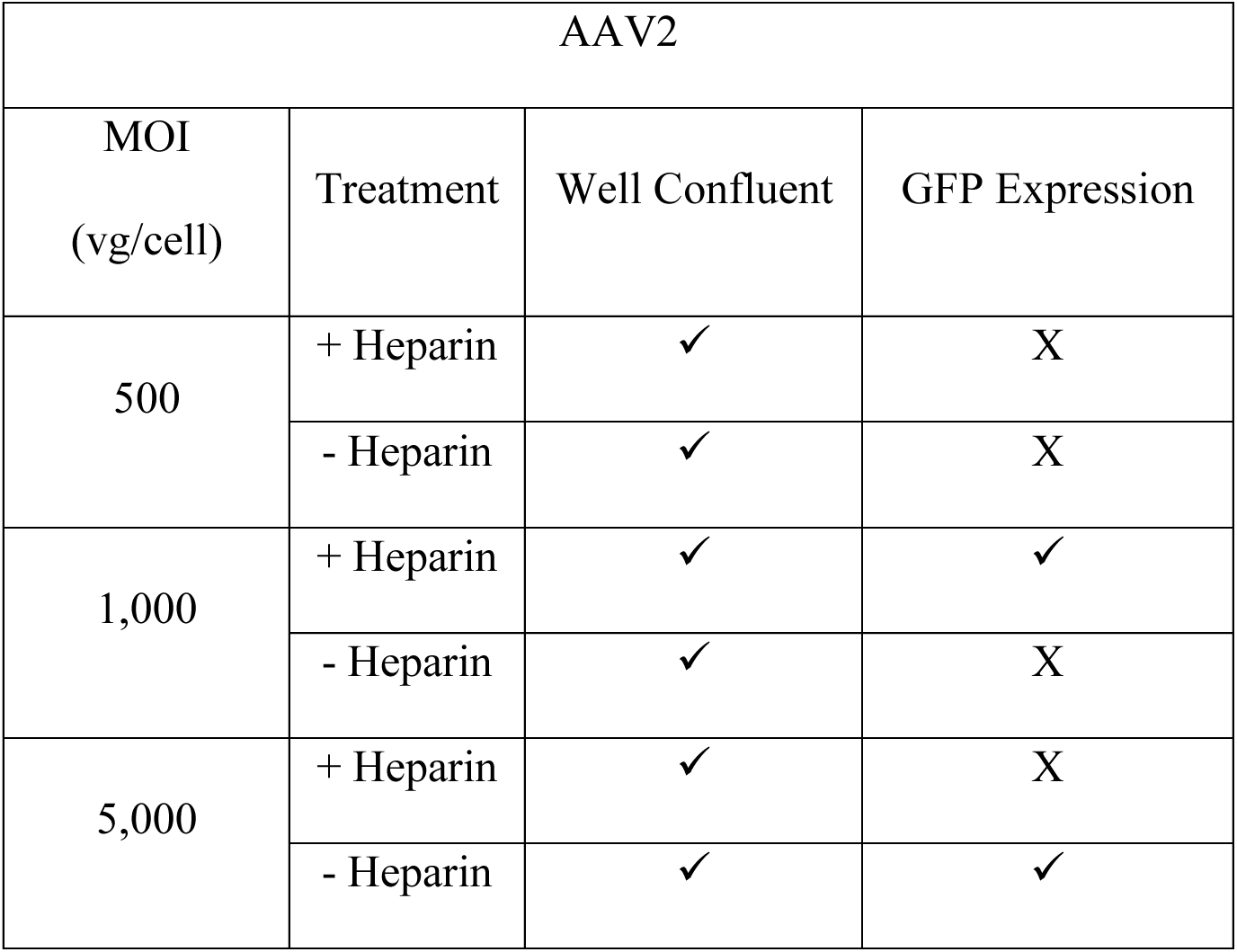
AAV2 MOI screen summarized results for well confluency and GFP expression. (✓) for well confluent represent no visual difference in confluency compared to the untreated control, with (X) having any visual difference. (✓) for GFP expression indicates visual confirmation of GFP expression when checked on an inverted microscope. All tested conditions displayed minimal virus-induced toxicity, but only cells transduced at a MOI of 5,000 without heparin visibly expressed GFP. Cultures were monitored up to 120 hours after infection.

| AAV2 |  |  |  |
| --- | --- | --- | --- |
| MOI<br>(vg/cell) | Treatment | Well Confluent | GFP Expression |
| 500 | + Heparin | ✓ | X |
|  | - Heparin | ✓ | X |
| 1,000 | + Heparin | ✓ | ✓ |
|  | - Heparin | ✓ | X |
| 5,000 | + Heparin | ✓ | X |
|  | - Heparin | ✓ | ✓ |

**Table S2.** AAV6 MOI screen summarized results for well confluency and GFP expression. (✓) for well confluent represent no visual difference in confluency compared to the untreated control, with (X) having any visual difference. (✓) for GFP expression indicates visual confirmation of GFP expression when checked on an inverted microscope. βTC cells visibly expressed GFP at 5,000 vg/cell irrespective of heparin, however at the cost of cell toxicity. Cultures were monitored up to 120 hours after infection.

| AAV6 |  |  |  |
| --- | --- | --- | --- |
| MOI<br>(vg/cell) | Treatment | Well Confluent | GFP Expression |
| 50 | Virus Only | ✓ | X |
|  | + Heparin | ✓ | X |
|  | + NM | ✓ | X |
|  | +Hep + NM | ✓ | X |
| 100 | Virus Only | ✓ | X |
|  | + Heparin | ✓ | X |
|  | + NM | ✓ | X |
|  | +Hep + NM | ✓ | X |
| 500 | Virus Only | ✓ | X |
|  | + Heparin | ✓ | X |
|  | + NM | ✓ | X |
|  | +Hep + NM | ✓ | X |
| 1,000 | Virus Only | ✓ | X |
|  | + Heparin | ✓ | X |
|  | + NM | X | X |
|  | +Hep + NM | X | X |
| 5,000 | Virus Only | X | ✓ |
|  | + Heparin | X | ✓ |
|  | +NM | X | X |
|  | +Hep + NM | X | X |

**Table S3.** AAV9 MOI screen summarized results for well confluency and GFP expression. (✓) for well confluent represent no visual difference in confluency compared to the untreated control, with (X) having any visual difference. (✓) for GFP expression indicates visual confirmation of GFP expression when checked on an inverted microscope. All virus-only wells displayed minimal cytotoxic effects and had GFP expression at 5,000 vg/cell and above. NM treatment of the cells appears to enhance transduction by AAV9 at low MOI, but simultaneously induces cell detachment. The level of NM-induced cell loss is proportional to the viral MOI added. Cultures were monitored up to 120 hours after infection.

| AAV9 |  |  |  |
| --- | --- | --- | --- |
| MOI<br>(vg/cell) | Treatment | Well<br>Confluent | GFP<br>Expression |
| 50 | - NM | ✓ | X |
|  | + NM | ✓ | X |
| 100 | - NM | ✓ | X |
|  | + NM | ✓ | ✓ |
| 500 | - NM | ✓ | X |
|  | + NM | X | ✓ |
| 1,000 | - NM | ✓ | ✓ |
|  | + NM | X | ✓ |
| 5,000 | - NM | ✓ | ✓ |
|  | + NM | X | X |
| 10,000 | - NM | ✓ | ✓ |
|  | + NM | X | X |

